# Immunoreactive peptide maps of SARS-CoV-2 and other human coronaviruses

**DOI:** 10.1101/2020.08.13.249953

**Authors:** Nischay Mishra, Xi Huang, Shreyas Joshi, Cheng Guo, James Ng, Riddhi Thakkar, Yongjian Wu, Xin Dong, Qianlin Li, Richard Pinapati, Eric Sullivan, Adrian Caciula, Rafal Tokarz, Thomas Briese, Jiahai Lu, W. Ian Lipkin

## Abstract

Serodiagnosis of SARS-CoV-2 infection is impeded by immunological cross-reactivity to the human coronaviruses (HCoV) SARS-CoV-2, SARS-CoV-1, MERS-CoV, OC43, 229E, HKU1, and NL63. Here we report the identification of humoral immune responses to SARS-CoV-2 and other HCoV peptides that can be used to detect asymptomatic, mild and, severe SARS-CoV-2 infections, and may enable the discovery of biomarkers for immunity following infection or vaccination.

## Main text

Differential serodiagnosis of human coronavirus (HCoV) exposure may be challenging due to cross-reactive immunity to SARS-CoV-2, SARS-CoV-1, MERS-CoV, OC43, 229E, HKU1, NL63^1-3^. Specific plaque reduction neutralization tests are labor-intensive, require work with live virus in high-level containment facilities, and detect only neutralizing antibodies. Here we report the use of proteome-wide high-density peptide microarrays to detect specific humoral immune responses to SARS-CoV-2 and other HCoV.

We established a microarray comprising >172,000 “12-amino acid (aa)” nonredundant peptides that tile the proteomes of known HCoVs with 11 amino acid overlap ^4-7^(Supplementary Table 1). Using these arrays, we examined the immunoreactivity of 132 plasma samples collected at two timepoints from fifty patients with active or recent SARS-CoV-2 infection, and 32 subjects including healthy controls, IgG positive for SARS-CoV-1, and exposure to other HCoVs (Supplementary Table 2 and 3). Subjects included 22 COVID-19 patients with severe illness (Group 1); 22 COVID-19 patients with mild illness (Group 2); 6 subjects with asymptomatic SARS-COV-2 infection (Group 3); 11 subjects with SARS-CoV-1 infection (Group 4); 10 healthy subjects (Group 5), and 11 subjects immunoreactive to other HCoV specific IgG ELISA. Plasma were heat inactivated, diluted, hybridized on HCoV peptide arrays, incubated with goat anti-human IgG and anti-human IgM antibodies, and scanned. Multidimensional scaling (MDS) was performed for IgM and IgG antibodies to differentiate peptides that were immunoreactive with COVID-19 patients (Groups 1-3) versus “non-COVID-19” control groups (Group 4-6). Details of samples, experimental design and data analysis are included in supplementary methods^4-7^.

IgM analysis revealed one linear epitope in the membrane glycoprotein (MADSNGTITVEELKKLLEQWN), which was reactive in 4/22 (19%) COVID-19 patients with mild disease and 8/22 (37%) patients with severe disease at late time point collection (Supplementary Figure 1 and Supplementary Table 4). IgG analysis revealed 37,237 peptides with statistically significant differences (p<0.05) in signal intensity between COVID-19 and control plasma. Multidimensional scaling (MDS) enabled separation of patients with COVID-19 (Groups 1-3) and controls (Groups 4-6) (Figure 1-A). 981 SARS-CoV-2 peptides with IgG signal intensity >10,000 arbitrary unity (AU) were used to assemble a heat map using R-Studio (Figure 1-B). Immunoreactive peptides included 566 from ORF1ab, 3 from ORF10, 243 from surface glycoprotein (S protein), 20 from ORF3a, 20 from membrane glycoprotein (mGP), 4 from ORF7a, 21 from ORF8, and 104 from nucleocapsid phosphoprotein (N). Peptides from “S” and “N” proteins had higher reactive intensity (higher AU) and rate of reactivity in comparison to peptides from other proteins (Figure 1-B). Immunoreactivity was higher in COVID-19 patients with severe vs mild disease, and in asymptomatic SARS CoV-2 infected subjects than in patients with mild disease. Samples from second timepoint collections were more reactive than first timepoint collections in severe, mild and asymptomatic SARS-CoV-2 infection (Figure 1-B). The presence of three continuous peptides that were reactive in samples from SARS-CoV-2 infection but not in control groups, were used to define 163 SARS-CoV-2 specific epitopes (Supplementary Table 5). Twenty-nine epitopes were selected based on sensitivity and specificity for SARS-CoV-2 infection status (Table 1). These included 11 epitopes (37.9%) in S protein (SP1-SP11), 8 (27.5%) epitopes in N protein (NP1-NP8), 6 (20.7%) epitopes in ORF1ab polyprotein (OP1-OP6), 2 (6.9%) in mGP protein (MP1 and MP2), one (3.4%) each from ORF3, and ORF8 proteins (Supplementary Figure 2). Table 1 indicates the location of each epitope on the SARS-CoV-2 proteome, its length, aa sequence, and the percentages of plasma samples that were immunoreactive in Groups 1-6. Immunoreactivity was higher in second time point plasma samples (Supplementary Figure 3) and in patients with more severe disease. In samples from patients with severe disease, 7 to 22 epitopes (mean of 13) were positive in second timepoint samples (24.8±6.8 days post onset of disease, POD) vs 0 to15 (mean of 7) epitopes in first timepoint samples (9.6±3.5 days POD). In patients with mild disease, 3 to 22 epitopes (mean of 8) were positive in second timepoint samples (34.7±8.3 POD) vs 0 to 12 epitopes (mean of 4) in first timepoint samples (12.9±5.9 POD). In asymptomatic subjects 1 to 9 epitopes (mean of 4) were positive at either time point (day of hospitalization and 14.5±4.6 days after the hospitalization). Plasma samples from 19 of 22 patients (86%) with mild disease were reactive with at least 1 of 29 epitope at the first time point; all 22 (100%) were reactive with at least 3 of 29 epitopes at the second time point. Plasma from 21 of 22 patients (95%) with severe disease were reactive with at least 1 epitope at first timepoint; all 22 (100%) were reactive with at least 6 epitopes at the second time point. All 6 (100%) asymptomatic cases were reactive with at least 2 of 29 epitopes in first and 3 of 29 in second timepoint collections.

**Table 1.**
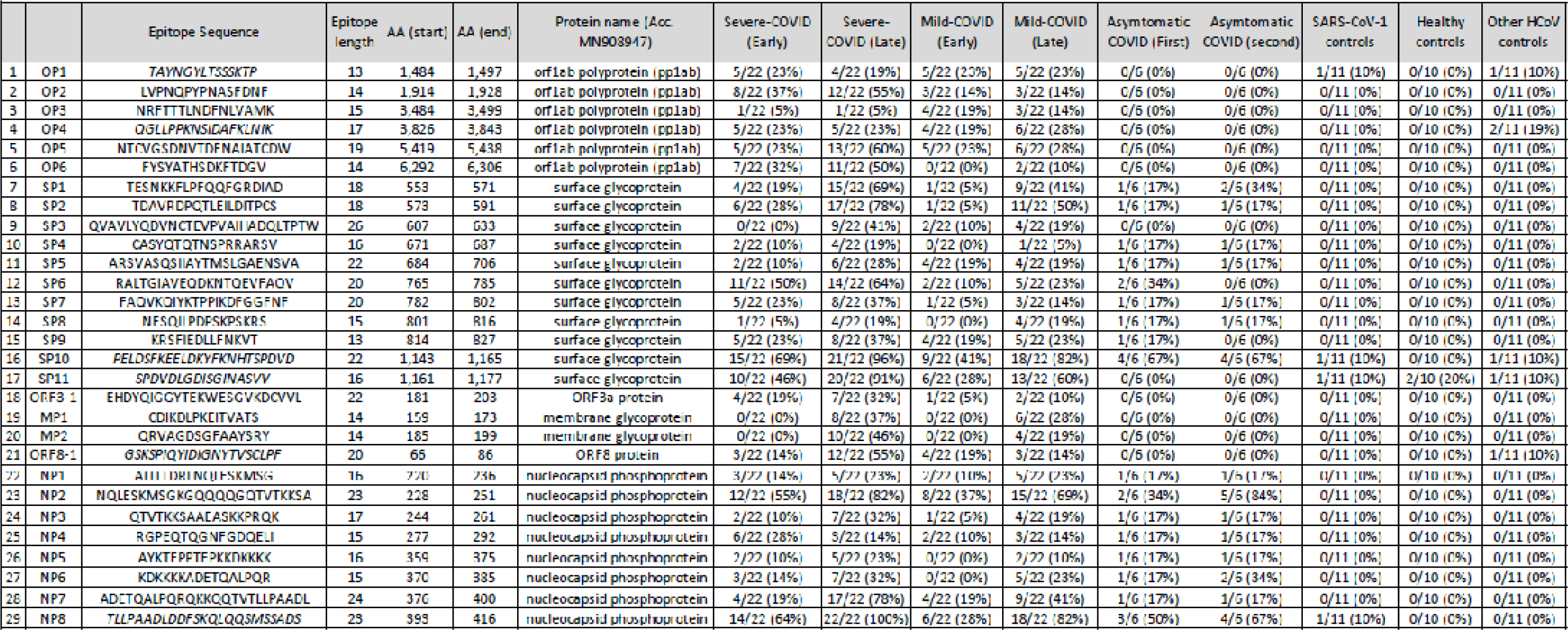
Characteristics of selected twenty-nine I&G linear epitopes for detection of SARS-CoV-2 infection Sequences (aa), length, aa location in protein is based on proteome of severe acute respiratory syndrome coronavirus 2 isolate Wuhan-Hu-1 (Accession no MN903947)

**Figure 1.**
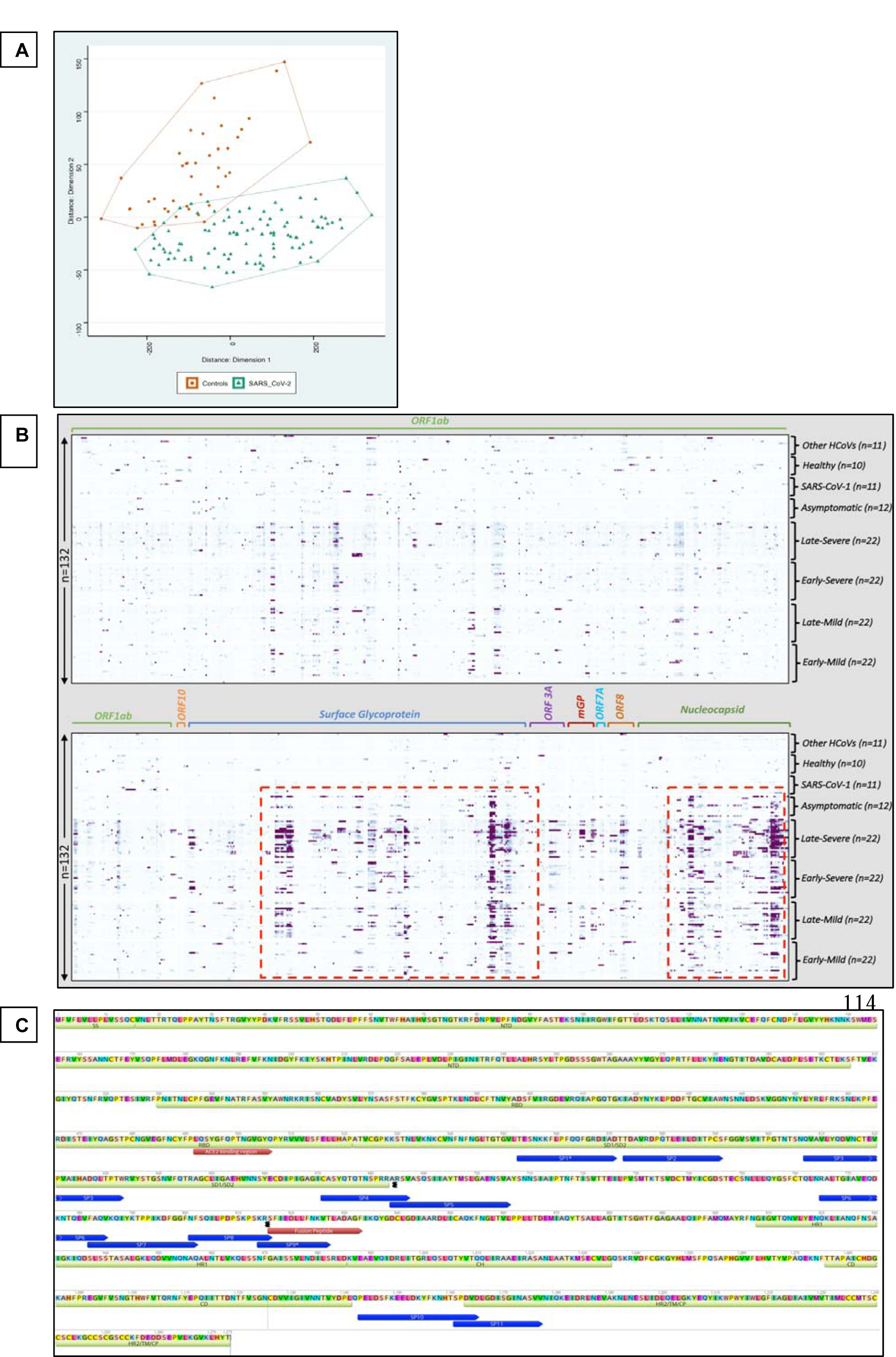
Discovery of immunoreactive IgG peptides for SARS-CoV-2. A] Multidimensional scaling (MDS) of differential IgG peptide signals in assays of sera from subjects with a history of infection with SARS-CoV-2 or without historical exposure to SARS-CoV-2 (controls). Based on MDS analysis, samples with exposure of SARS-CoV-2 samples (green) versus controls (red) clustered in two separate groups. B] Proteome wide linear epitope mapping of SARS-CoV-2 specific IgG antibodies by a HCoV peptide array. X axis represents 981 peptides from SARS-CoV-2 proteins, Y axis represents 132 samples tested using the HCoV peptide array. Heatmap is plotted with individual peptide intensity in AU for each of the 132 plasma samples. Panel grids show highly reactive areas in S and N proteins. * ORF1ab protein is large so divided partly in lower panel due to larger size. C] Location of Spike epitopes (SP1-SP11) is shown that in dark blue arrows. Primary structure domains are colored by green bars: SS, signal sequence; S2′, S2′ protease cleavage site; FP, fusion peptide; HR1, heptad repeat 1; CH, central helix; CD, connector domain; HR2, heptad repeat 2; TM, transmembrane domain; CT, cytoplasmic tail. “{” symbols indicate protease cleavage sites. Brown arrows denote ACE2 binding region and fusion peptide regions. *Epitopes with neutralizing antibody binding potential (SP1*, and SP9*).

Through use of plasma from subjects with known exposure to other HCoV we identified specific epitopes in HKU1 (n=15), NL63 (n=10), OC43 (n=14), 229E (n=5), and SARS-CoV-1 (n=9) (Supplementary Table 6). We were unable to identify MERS-specific epitopes due to lack of cognate sera. More than 90% of samples from patients exposed to SARS-CoV-2 and controls showed a wide range of reactivity to epitopes from seasonal coronaviruses (HKU1, NL63, OC43 and 229E). A Pre-COVID study from Beijing, China showed that large proportions (>70%) of healthy children and adults have evidence of anti-S IgG antibodies against these four HCoVs^8^. The immunoreactivity of SARS-CoV-1 epitopes to SARS-CoV-2 infected patients can be explained by cross reactivity due to proteome homology ^2,3^, and immunoreactivity of SARS-CoV-1 epitopes in healthy controls maybe due to historical unknown exposure during SARS-2003 epidemic in Guangdong province ^9,10^.

The 29 immunoreactive epitopes were mapped to the proteome of SARS-CoV-2 (acc. number MN908947) (Supplementary Figure 2). ORF1ab epitopes (OP1-OP6) were dispersed throughout the protein. Eleven linear epitopes from “S” protein (SP1-SP11) were mapped outside of the RBD (Receptor Binding Domain) (Figure 1-C). Four SP1-SP4 epitopes were located in the SD1/SD2. Four SP5-SP8 epitopes were located between SD1/SD2 and fusion peptide region. The SP9 epitope overlapped the Fusion Peptide (FP). The SP10 epitope is located between CD (Connector Domain) region and HR2 (Heptad Repeat 2). The SP11 epitope was located at the beginning of the HR2 region of the S2 subunit of the surface glycoprotein (Figure 1-C). The SP1, and SP9 peptides identified in this study were recently reported as potentially linked to neutralization^11^. Ten of eleven spike epitopes (SP1-SP10) were located in regions of high immunoreactivity in a recent pre-print from Li et al^12^. The SP10 epitope was the most sensitive diagnostic spike epitope in subjects with severe (69% first time point, 96% second timepoint), mild (41% first timepoint, 82% second timepoint), and asymptomatic SARS-CoV-2 (67% either timepoint) infections. The NP2 and NP8 epitopes were the most sensitive “N” protein epitopes. In severe disease NP2 immunoreactivity was found in 55% of subjects at the first timepoint, and 82% at the second timepoint. NP8 immunoreactivity was found in 64% of subjects at first timepoint, and 100% at second time point. In mild disease, NP2 immunoreactivity was found in 37% of subjects at the first timepoint, and 69% at the second timepoint. NP8 immunoreactivity was found in 28% of subjects at the first timepoint, and 82% at the second timepoint. In asymptomatic cases, immunoreactivity was 59% for either NP2 or NP8. In severe disease SP11 immunoreactivity was found in 46% of subjects at the first time point, and 91% at the second time point. In mild disease SP11 immunoreactivity was found in 28% of subjects at the first time point, and 60% at the second time point. No asymptomatic cases were immunoreactive with SP11. Epitopes for ORF1ab (OP1-OP6) showed reactivity in cases with mild and severe disease at both timepoints but not in asymptomatic infections. Only second timepoint samples from severe and mild diseases were immunoreactive to epitopes for mGP, MP1 and MP2 (37% and 26% for severe disease and 28% and 19% for mild disease)

The HCoV peptide array enabled serological diagnosis of SARS-CoV-2 and other coronavirus infections. Immunoreactivity profiles differed with severity of illness and over the timecourse of infection. The sensitivity of the array with this sample set was 92% at ∼10-13 days POD, 100% at ∼24-35 days POD with severe and mild COVID-19. Assays were positive in all asymptomatic, hospitalized subjects; however, only six subjects were examined and the duration of infection is unknown. The HCoV array platform is too complex and expensive for routine clinical microbiology. However, the peptides defined here can be transferred to a wide range of platforms including microarrays, ELISA, RIA, lateral flow, western blot, and bead-based assays, where they may facilitate diagnostics, epidemiology, and vaccinology.

## Methods

### Human coronavirus (HCoV) peptide array design

We designed a programmable peptide microarray that can accommodate up to 3 million distinct linear peptides on a 75mm x 26mm slide. The array can also be divided into 12 sub arrays, each containing approximately ∼173,000 “12-mer peptides” (Nimble Therapeutics Inc, WI, USA). The 12-mer format is based on the observation that serum antibodies bind linear peptide sequences ranging from 5 to 9 amino acid (aa) and bind most efficiently when targets are flanked by additional aa^13^. To enable differential detection of antibodies specific for SARS-CoV-2 infections, we created a database comprising the proteomes of seven human coronaviruses: SARS-CoV-2, SARS, MERS, NL-63, OC-43, 229E and HKU1 (Supplementary Table 1). We also included two bat coronavirus proteomes similar to SARS-CoV-2^14^. 1000 randomly selected 12 aa long scrambled peptides were added for background correction and nonspecific binding of peptides. For each virus selected, we downloaded all available protein sequences available before January 2020 from the NCBI and Virus Pathogen Database and Analysis Resource (VIPR) protein databases. We then created a peptide database comprising overlapping 12-mer peptides that tiled the whole proteome of each of these agents with 11 amino acid (aa) overlap in a sliding window pattern ^4-7^. The selected peptide sequences were passed through a redundancy filter to yield 283630 unique peptide sequences and 24779921 non-unique (present in more than one virus) for a total of 172,665 peptides. Redundant peptides were excluded prior to synthesis. The individual peptides in the library were printed in random positions on the peptide array to minimize the impact of locational bias.

### Samples & experimental design

The study was approved by the Medical Ethical Committee of Sun Yat-Sen University (approval number 2020-060). An informed and written consent was obtained by patients. A total 132 plasma samples were tested and analyzed using HCoV peptide arrays (Supplementary Table 2). Samples were divided into six groups: group 1] COVID-19 patients with non-severe (mild) disease (n=22); group 2] COVID-19 patients with severe disease (n=22); group 3] patients with SARS-CoV-2 infections but no-symptoms (asymptomatic COVID-19) (n=6); group 4] SARS-2003 IgG positive cases (n=11); group 5] other banked HCoV IgG positive controls (n=11) (Supplementary Table 3); and group 6] healthy controls (n=10). The average age was 44.0±16.73 years for the mild COVID-19 group, 60.1±12.37 years for the severe COVID-19 group, and 43.5±15.08 years for the asymptomatic COVID-19 group. The average age for SARS-CoV-1 IgG positive control group was 24.4±5.0 years. Plasma samples were collected at two different time points, a minimum of two weeks apart, from group COVID-19 patients (groups 1, 2 and 3). The first time point (early) was collected at 12.9±5.9 post onset of disease (POD) for the mild disease group, and at 9.6±3.5 days POD for the severe disease group. The second time point (late was collected at 34.7±8.3 days POD for the mild disease group, and at 24.8±6.8 days POD for the severe disease group. For asymptomatic group, the first time point was collected on the day of hospitalization; the second time point was collected at 14.5±4.6 days after the day of hospitalization. Non COVID-19 samples from control groups (group 4, group 5 and group 6), were from adults without any evidence or history of infection with SARS-CoV-2. All COVID-19 patients were tested for SARS-CoV-2 RNA in respiratory specimens using the China FDA approved Novel Coronavirus (2019-nCoV) Real Time RT-PCR kit from LifeRiver Ltd. (Catalog #: RR-0479-02) real-time RT-PCR^15^. The diagnosis of COVID-19 pneumonia, and severity criteria were assessed at Guangdong CDC based on the New Coronavirus Pneumonia Prevention and Control Program (6th edition) published by the National Health Commission of China^15^. Other clinical information, comorbidities, symptoms and treatment for COVID-19 for mild, severe, and asymptomatic cases are presented in Supplementary Table 2.

### HCOV peptide array synthesis, sample binding and processing

Peptide synthesis was accomplished by light-directed array synthesis in a Nimble therapeutics Maskless Array Synthesizer (MAS) using an amino-functionalized substrate coupled with 6-amino hexanoic acid as a spacer and aa derivatives carrying a photosensitive 2-(2-Nitrophenyl) propyl-oxy-carbonyl-group (NPPOC). Coupling of amino acids was done using pre-activated amino acid with activator (HOBT/HBTU) and Ethyl-di-iso-propylamine in DMF for 5–7 minutes before flushing the substrate. Cycles of coupling were repeated until 12-mer peptides were synthesized. Intermediate washes on the arrays were done with N-Methyl-2-pyrrolidone (NMP) and site-specific cleavage of the NPPOC group was accomplished by irradiation of an image created by a Digital Micro-Mirror Device (Texas Instruments, SXGA+ graphics format), projecting light with a 365-nm wavelength. Final de-protection to cleave off the side-chain protecting groups of the amino acids was done with Trifluoroacetic acid (TFA)/Water/ Trisopropylsilane for 30 minutes.

Before loading, plasma samples were heat inactivated at 56^0^ C for 30 minutes. Plasma samples were diluted (1:50) with binding buffer (0.1M Tris-Cl, 1% alkali soluble casein, 0.05% Tween-20, and water). The peptide arrays were incubated overnight at 4°C on a flat surface with individual sample/sub-array. Overnight sample incubation was followed by three 10-minute washes with 1X TBS-T (0.05% Tween-20) at room temperature (RT). Secondary antibodies IgG (cat no. 109-605-098, Alexa Fluor 647-AffiniPure Goat Anti-Human IgG, Fcy fragment specific, Jackson ImmunoResearch Labs) and IgM (cat no. 109-165-129, Cy™3 AffiniPure Goat Anti-Human IgM, Jackson ImmunoResearch Labs) were diluted in 1X PBS at a concentration of 0.1µg/ml, and arrays were incubated in Plastic Coplin Jar (cat no. S90130, Fisher Scientific) for 3 hours at RT with gentle shaking. Secondary antibody incubation was also followed by three 10-minute washes with 1X TBST at RT. After a final wash, the arrays were dried and scanned on a microarray scanner at 2-μm resolution, with an excitation wavelength of 635 nm (IgG) and 532 nm (IgM). The images were analyzed using the PepArray analysis program. The fluorescent signals were converted into arbitrary unit (AU) intensity plots ranging minimum to maximum intensity 0-65,000 AU.

### Experimental Design and data analysis

We employed dilutions of 1:50 and called a peptide signal positive if it was above the threshold (mean ±2 SD readings of random peptides, >10,000 Arbitrary unit (AU) for IgG and IgM analysis). A cut-off threshold for peptide recognition was defined as mean ±2 times the standard deviation (SD) of the mean intensity value of all negative controls^16^. The normalization, background correction and statistical comparison of peptide microarray intensities between groups was performed using the edgeR package^17^.

### Data analysis for IgG and IgM

Fold changes and standard errors were estimated by fitting a linear model for signal intensities generated by each peptide, applying empirical Bayesian smoothing to the standard errors, and then determining those peptides that yielded statistically significant signal by contrasting linear models for each peptide between SARS-CoV2 and Control samples at a significance value of <0.05^18^. The analysis was performed to differentiate peptides that were immunoreactive with COVID-19 patients (groups 1, 2 and 3) and versus control groups (groups 4, 5 and 6) samples. A multidimensional scaling (MDS) plot was generated using signal data for these significant peptides. Signal data points were filtered such that only peptides that showed signal >10,000 AU in any sample were retained. For IgG, the >10,000 AU filtration step reduced the initial number of peptides from 172665 to 79714 for further analysis. A total of 37,237 peptides (18533 from the COVID-19 group and 18704 from the control group) yielded group-specific differences (p<0.05) in signal intensity. Multidimensional scaling (MDS) demonstrated the capacity of these peptides to separate samples collected from patients with COVID-19 and controls (Figure 1-A). For IgM analysis, >10,000 AU filtration step reduced the initial number of peptides from 172665 to 24728 for further analysis. A total of 10,816 peptides (7144 from the COVID-19 group and 3672 from the control group) peptides yielded group-specific differences (p<0.05) in signal intensity. Multidimensional scaling (MDS) efforts did not separate samples collected from patients with COVID-19 and controls (Supplementary Figure 1). However, the presence of three continuous peptides that were reactive in samples from SARS-CoV-2 infected subjects (irrespective of disease status) but not in control groups, allowed the identification of 16 SARS-CoV-2 specific IgM epitopes (supplementary table 4). The code for reassembly and plots was prepared using Rstudio v 1.2.5019^19^. The Heatmap plots were generated using ggplot2 package^20^. A custom color-blind friendly color pallete was used to make the plots. Alignment of reactive epitopes on SARS-CoV-2 proteome was performed using Geneious version 10.0.9.

## Author contributions

NM, XH, JL and WIL contributed to study design, data interpretation and manuscript preparation. YW, XD, QL, SJ, and CG contributed to data acquisition and graphics. TB, and RaT had roles in data interpretation. JN and RiT had roles in data generation. All authors reviewed and approved the final version of the manuscript.

## Acknowledgements

We thank Lokendra V Chauhan and Teresa Tagliafierro for technical assistance. Work at the Columbia University was supported by the Chau Hoi Shuen Foundation and the Marin Community Foundation. Work at Sun Yat-Sen was funded by the Guangdong Scientific and Technological Research for COVID-19 (202020012612200001) and the National Science and Technology Major Project (No. 2018ZX10101002-001-001). We thank Nimble Therapeutics Team (Jigar Patel, Brad Garcia and Daniel Agnew) for array synthesis and technical support.

## Supplementary Figures and tables

**Supplementary Figure 1.**
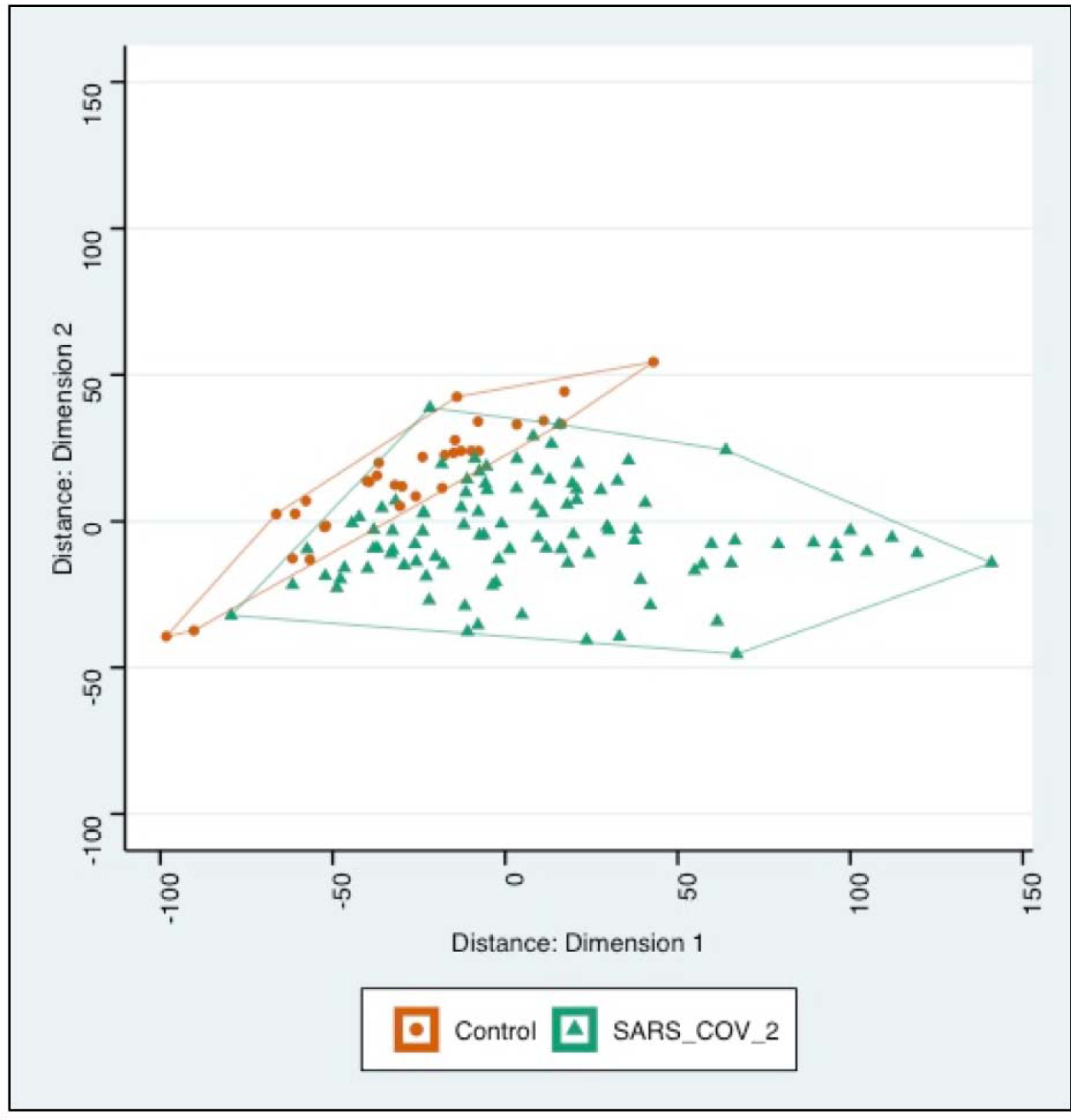
Multidimensional scaling (MDS) of differential IgM peptide signals in assays of sera from subjects with a history of infection with SARS-CoV-2 or without historical exposure to SARS-CoV-2 (controls). Based on MDS analysis, samples with exposure of SARS-CoV-2 samples (green) versus controls (red) clustered in overlapping groups.

**Supplementary Figure 2.**
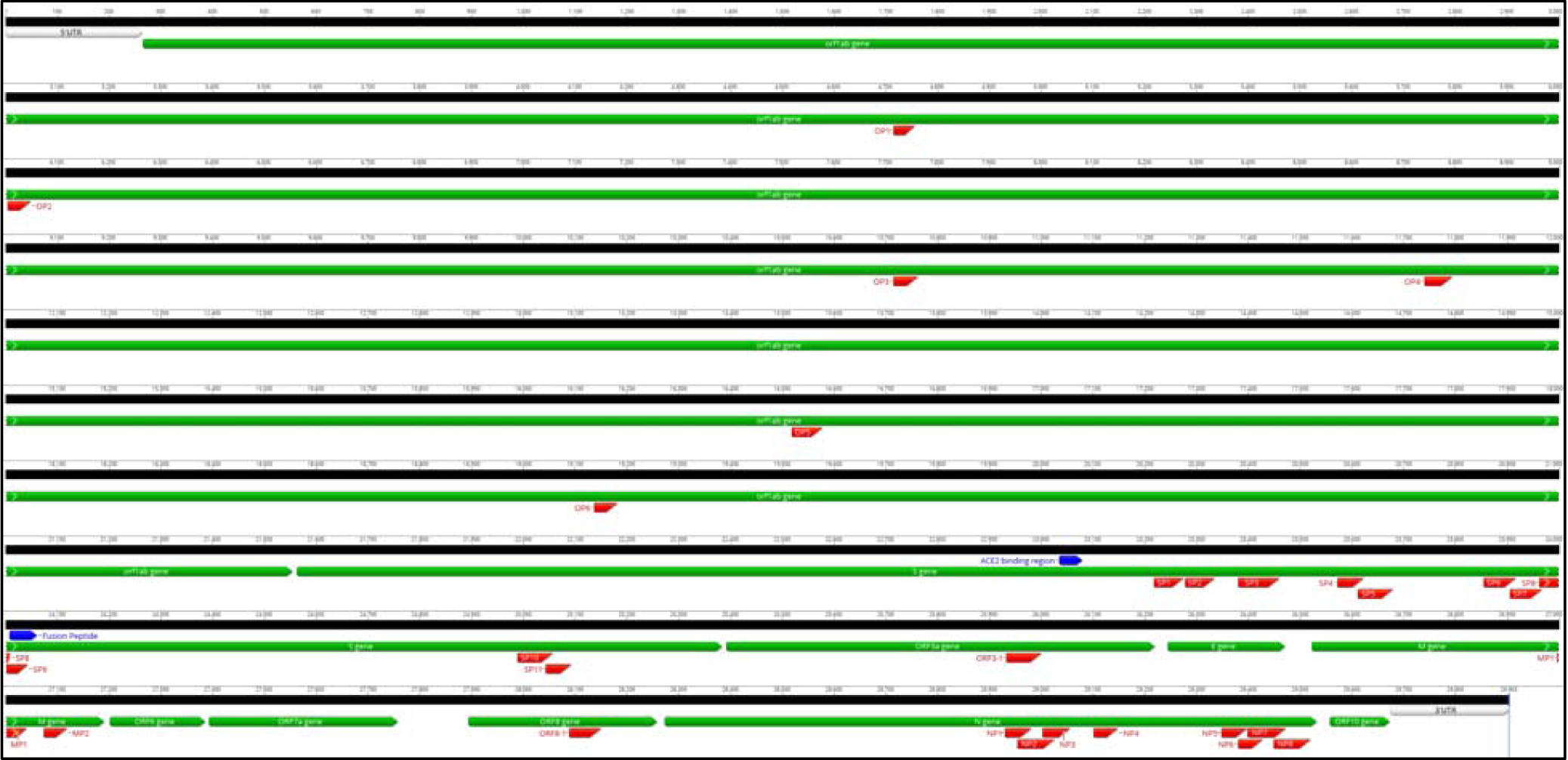
Notation of twenty-nine immunogenic B cell IgG epitopes on the proteome of SARS-CoV-2 (Accession number MN908947)

**Supplementary figure 3.**
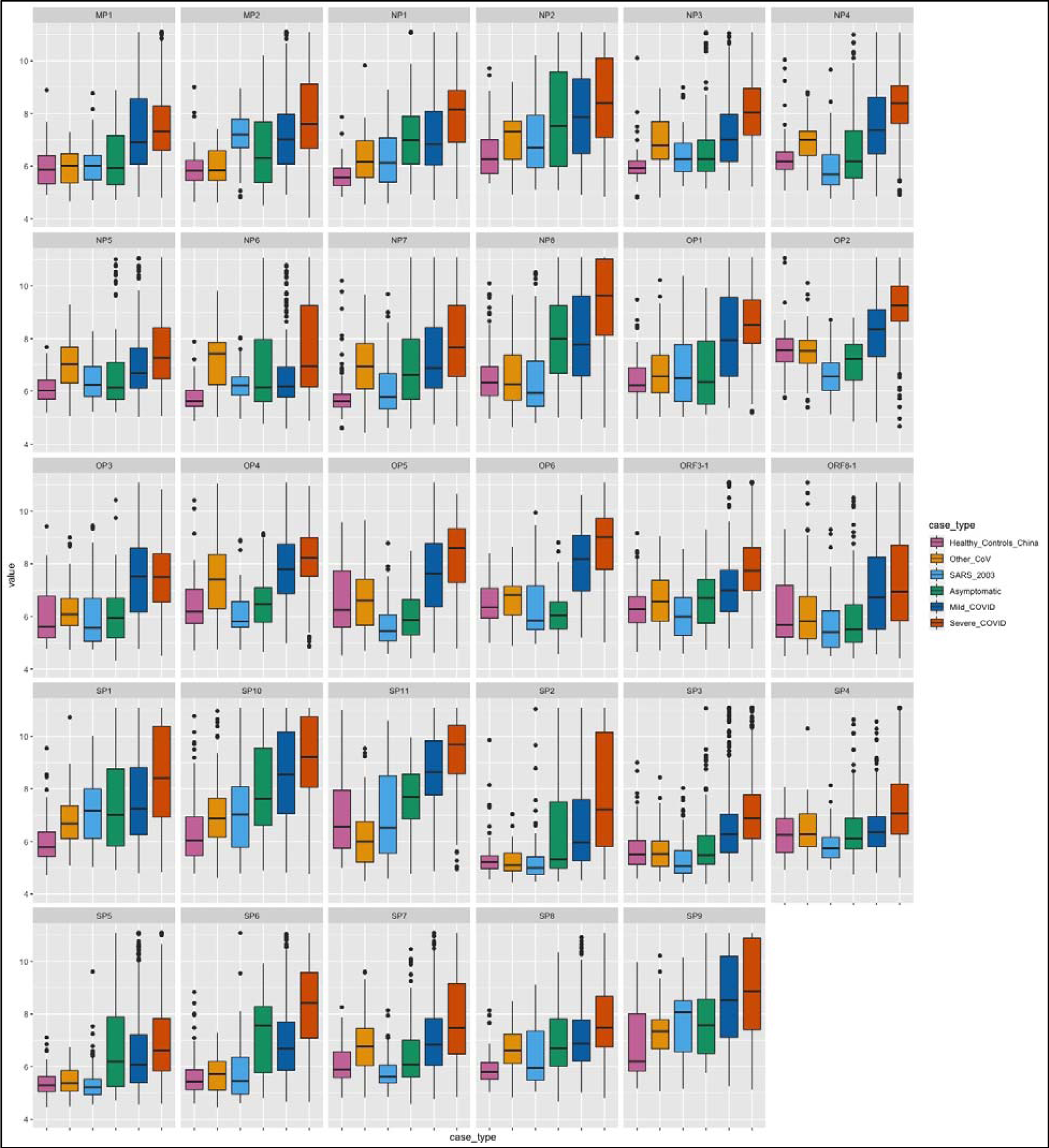
Reactivity intensities of each peptide from 29 immunogenic linear IgG epitopes are plotted on log scale (y-axis). Reactivity to selected peptides is plotted for corresponding groups shown in different colors (x-axis).

**Supplementary Table 1.** Design and characteristics of HCoV-peptide array.

|  | Virus database | No. of protein sequences | Total redundant "12-aa" peptides | Total unique "12-aa" peptides |
| --- | --- | --- | --- | --- |
| 1 | *SARS-CoV-2-MN908947-CDS* | 10 | 9591 | 9591 |
| 2 | *Human-229E-CoV-PDB* | 662 | 576999 | 15756 |
| 3 | *Human-HKU1-CoV-PDB* | 721 | 554447 | 16390 |
| 4 | *Human-NC63-CoV-PDB* | 1320 | 1000287 | 16392 |
| 5 | *Human-OC43-CoV-PDB* | 2667 | 2679272 | 29635 |
| 6 | *MERS-CoV-PDB* | 7253 | 8604059 | 54188 |
| 7 | *SARS-CoV-PDB* | 1996 | 1484140 | 29791 |
| 8 | *Human-CoV-variants-VIPR-PDB* | 12131 | 9880717 | 101145 |
| 9 | *Bat-CoV-ZC45* | 11 | 9678 | 9678 |
| 10 | *Bat-CoV-ZXC21* | 11 | 9655 | 9655 |
| 11 | *Redundant-non-specific-peptides* | NA | NA | 1000 |
| Total number of unique and redundant peptides |  |  | 24779921 | 283630 |
| **Total peptides in HCoV-SeroChip** |  |  | **172665** |  |

**Supplementary Table 2.** Clinical characteristics of patients by groups with SARS-CoV-2 infection and controls used in this study.

|  | Asymptomatic COVID<br>(n=6) | Mild COVID<br>(n=22) | Severe COVID<br>(n=22) | SARS-CoV-1<br>(n=11) | Healthy<br>(n=10) | Other HCoV<br>(n=11) |
| --- | --- | --- | --- | --- | --- | --- |
| Gender (M/F) | 2/4 | 13/9 | 14/8 | 0/11 | -- | -- |
| Age (years) | 43.5±15.08 | 44.0±16.73 | 60.1±12.37 | 24.4±5.0 | -- | -- |
| Time of collection (post onset of symptoms days, POD) |  |  |  |  |  |  |
| First collection | day of hospitalisation | 12.9±5.9 | 9.6±3.5 | -- | -- | -- |
| Second collection | 14.5±4.6 | 34.7±8.3 | 24.8±6.8 | -- | -- | -- |
| Symptoms |  |  |  |  |  |  |
| Cough | 0(0%) | 19(86%) | 15(68%) | -- | -- | -- |
| Fever | 1(17%) | 16(73%) | 19(86%) | -- | -- | -- |
| Dyspnea | 0(0%) | 2(9%) | 5(23%) | -- | -- | -- |
| Snivel | 0(0%) | 3(14%) | 1(5%) | -- | -- | -- |
| Sore throat | 0(0%) | 4(18%) | 5(23%) | -- | -- | -- |
| Diarrhea/Vomit/Stomach ache | 0(0%) | 7(32%) | 8(36%) | -- | -- | -- |
| Underlying diseases |  |  |  |  |  |  |
| Cancer | 1(17%) | 2(9%) | 0(0%) | -- | -- | -- |
| Thyroid disease | 1(17%) | 0(0%) | 2(9%) | -- | -- | -- |
| Diabetes | 0(0%) | 2(9%) | 3(14%) | -- | -- | -- |
| Hypertension | 0(0%) | 5(23%) | 6(27%) | -- | -- | -- |
| Arthritis | 0(0%) | 1(5%) | 0(0%) | -- | -- | -- |
| Kidney disease | 0(0%) | 1(5%) | 1(5%) | -- | -- | -- |
| Appendicitis History | 0(0%) | 1(5%) | 1(5%) | -- | -- | -- |
| Gastritis history | 0(0%) | 0(0%) | 1(5%) | -- | -- | -- |
| Spinal diseases | 0(0%) | 0(0%) | 2(9%) | -- | -- | -- |
| Bronchiectasis | 0(0%) | 0(0%) | 1(5%) | -- | -- | -- |
| Skin disease | 0(0%) | 0(0%) | 1(5%) | -- | -- | -- |
| Treatment |  |  |  |  |  |  |
| Antibiotic | 3(50%) | 17(77%) | 22(100%) | -- | -- | -- |
| Antivirus | 5(83%) | 19(86%) | 21(95%) | -- | -- | -- |
| Interferon | 0(0%) | 1(5%) | 3(14%) | -- | -- | -- |
| TCM | 1(17%) | 7(32%) | 6(27%) | -- | -- | -- |
| Globulin | 0(0%) | 11(50%) | 21(95%) | -- | -- | -- |
| Chloroquine | 5(83%) | 16(73%) | 9(41%) | -- | -- | -- |

**Supplementary Table 3.**
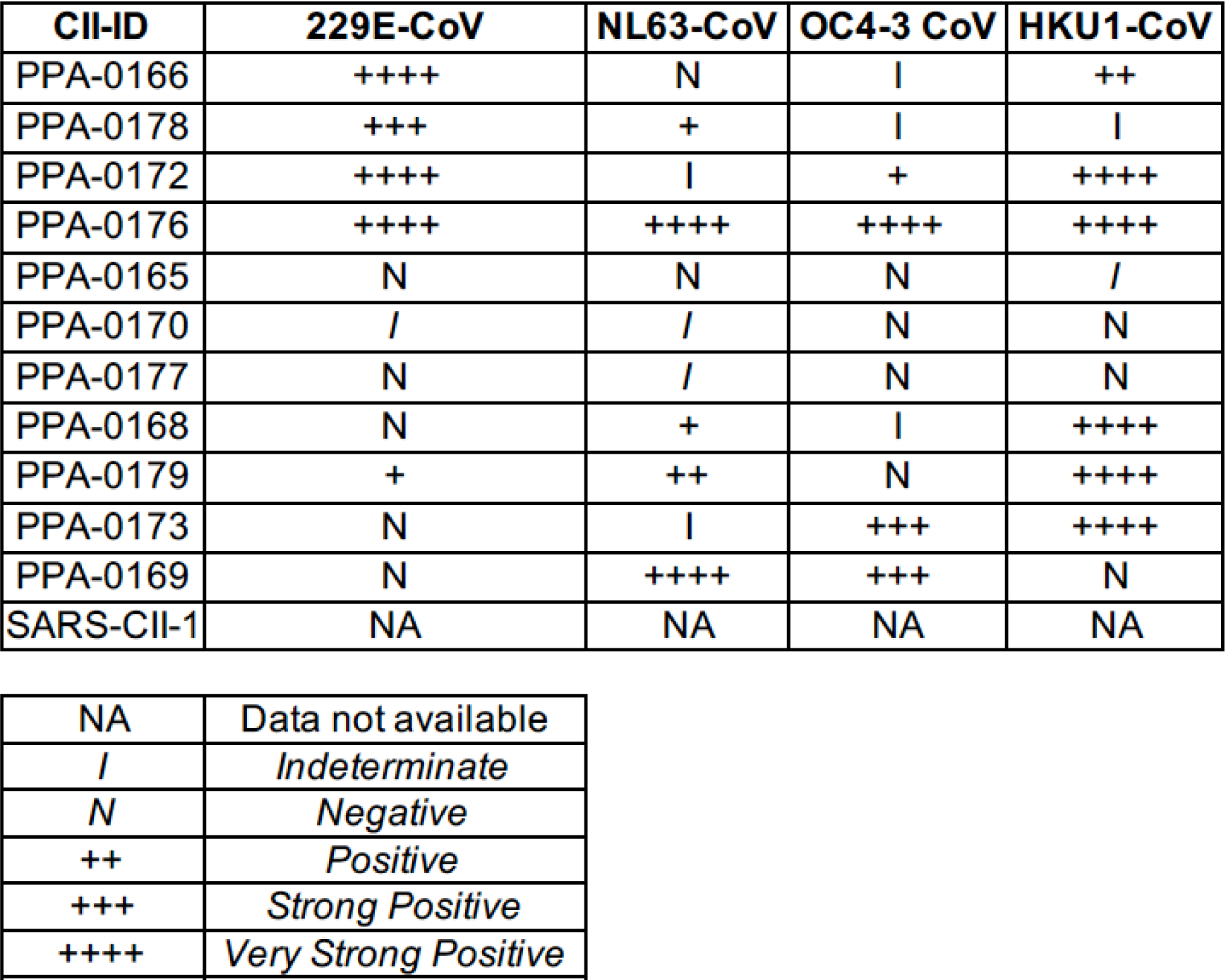
IgG Immunoreactivity of other HCoV controls using S protein based specific ELISAs for four seasonal human coronaviruses (OC43, 229E, HKU1 and NL63).

| CII-ID | 229E-CoV | NL63-CoV | OC4-3 CoV | HKU1-CoV |
| --- | --- | --- | --- | --- |
| PPA-0166 | ++++ | N | I | ++ |
| PPA-0178 | +++ | + | I | I |
| PPA-0172 | ++++ | I | + | ++++ |
| PPA-0176 | ++++ | ++++ | ++++ | ++++ |
| PPA-0165 | N | N | N | I |
| PPA-0170 | I | I | N | N |
| PPA-0177 | N | I | N | N |
| PPA-0168 | N | + | I | ++++ |
| PPA-0179 | + | ++ | N | ++++ |
| PPA-0173 | N | I | +++ | ++++ |
| PPA-0169 | N | ++++ | +++ | N |
| SARS-CII-1 | NA | NA | NA | NA |

|  |  |
| --- | --- |
| NA | Data not available |
| I | *Indeterminate* |
| N | *Negative* |
| ++ | *Positive* |
| +++ | *Strong Positive* |
| ++++ | *Very Strong Positive* |

**Supplementary Table 4.**
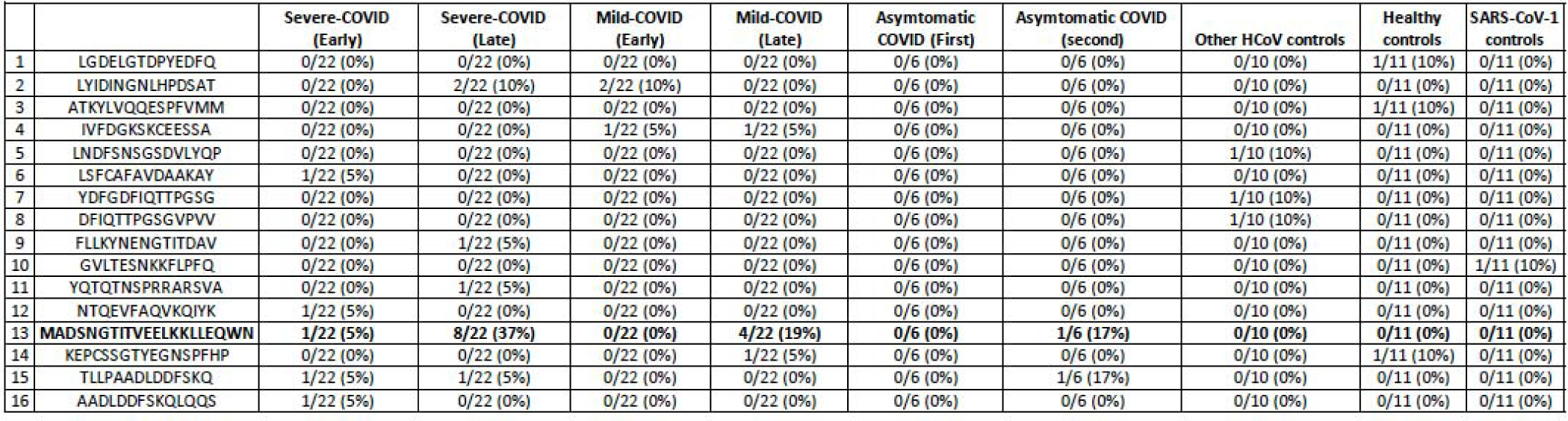
Characteristics of selected sixteen IgM linear epitopes for detection of SARS-CoV-2 infection. Sequences (aa) are based on proteome of Severe acute respiratory syndrome coronavirus 2 isolate Wuhan-Hu-1 (Accession no. MN908947).

**Supplementary Table 5.**
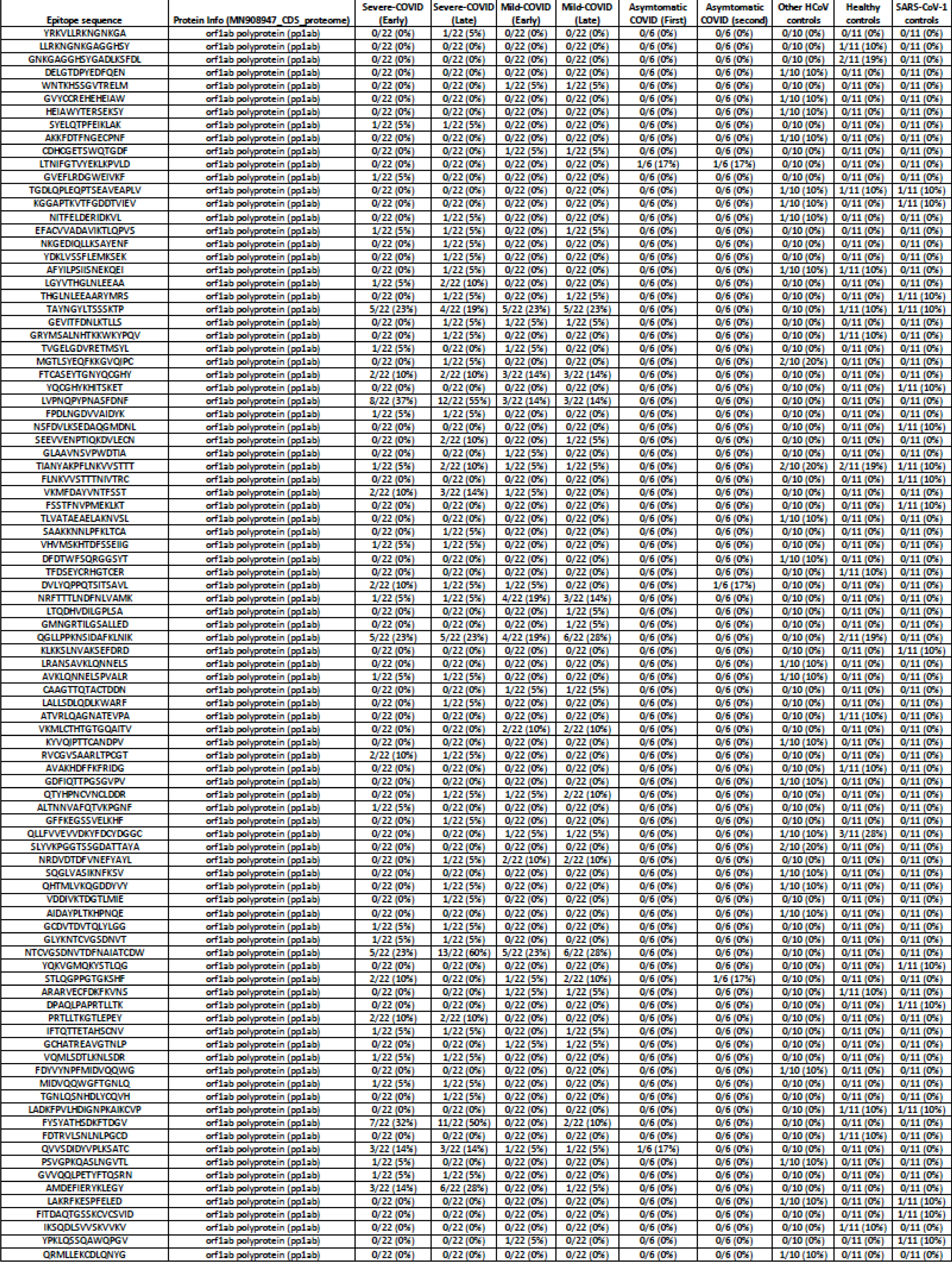
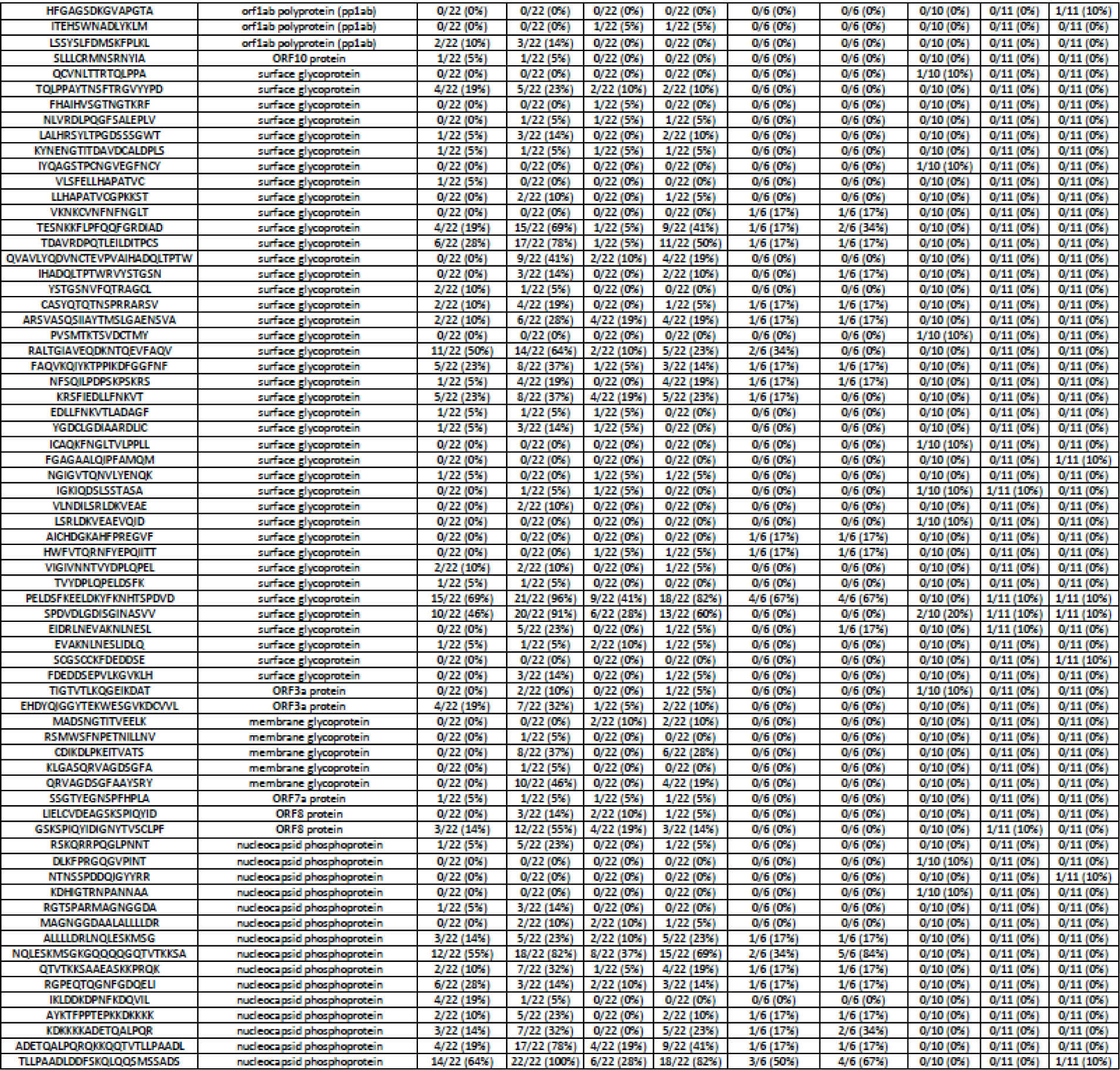
List of all 163 IgG linear epitopes for detection of SARS-CoV-2 infection. Sequences (aa) are based on proteome of Severe acute respiratory syndrome coronavirus 2 isolate Wuhan-Hu-1 (Accession no. MN908947)

**Supplementary table 6.**
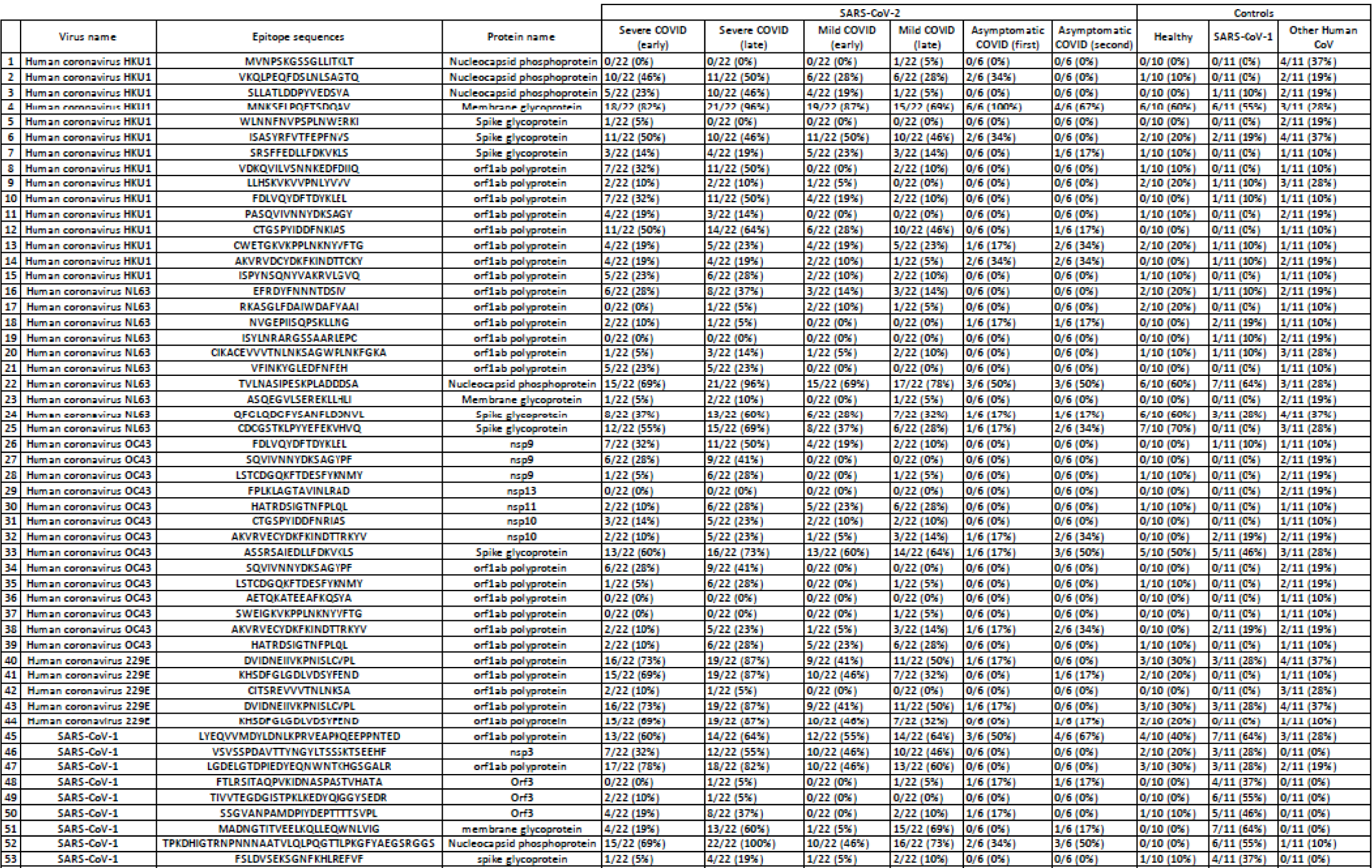
List of selected 55 IgG linear IgG epitopes for detection of HKU-1, NL63, OC43, 229E, SARS-CoV-1. Sequences (aa) are based on proteome of reference sequences from NCBI.

## Notes

https://github.com/ciibioinformatics/COVID19_publication

### Competing Interest Statement

NM, SJ, CG, RaT, TB, and WIL are listed as the inventors on a patent filed that is related to findings in this study.
Application: 63/005,865
Title: Peptide Sequences for Detection and Differentiation of Antibody Responses to SARS-CoV-2 and Other Human Corona Viruses
Application Type PRO - Provisional
Status: Filed
Country: United States
IRs: CU20309
IR Received: 2020-04-02
Filing Date: 2020-04-06
Inventor(s) : Nischay Mishra; W. Ian Lipkin M.D.; Rafal Tokarz; Shreyas Joshi; Cheng Guo; Thomas Briese;

